# Single-step monolithic chromatography efficiently purifies diverse Pseudomonas aeruginosa phages with therapeutic-grade endotoxin reduction

**DOI:** 10.1101/2025.02.03.636326

**Authors:** Arne Echterhof, Tejas Dharmaraj, Patrick Blankenberg, Maryam Hajfathalian, Francis Blankenberg, Jessica C. Sacher, Paul L. Bollyky

## Abstract

Bacteriophages are increasingly explored as antibacterial agents for therapeutic applications and pathogen biocontrol. Rigorous quality control and removal of microbial growth byproducts is essential for safety and compliance with regulatory standards for the use of phages; for phages isolated from Gram-negative bacterial lysates, this involves removal of endotoxin (lipopolysaccharide/LPS) and contaminating host proteins. Here, we describe the development of CIMmultus OH-Monolithic chromatography as a single-step purification process for five tailed anti-pseudomonal phages, including myoviral and siphoviral morphologies, as well as so-called “jumbo” phages, yielding therapeutically compliant samples suitable for human IV administration. Using preferential exclusion chromatography with a potassium phosphate gradient, we achieved near-complete removal of endotoxin (99.96-100.00% depletion) and protein impurities (95.8-99.7% depletion). We found that phage recovery post-purification was inversely associated with tail length and hydrodynamic diameter, with shorter-tailed and smaller phages demonstrating greater recovery. We conclude that single-step CIMmultus OH-monolithic chromatography is scalable, reproducible, and free from hazardous chemicals, supporting its integration into industrial phage purification processes.

## Introduction

Bacteriophages (phages) are viruses that infect bacteria^1^. Given their selective host range^2^, potent antibacterial activity^3^, and role in averting antibiotic overuse^4,5^, phages are increasingly pursued as antibacterial options—particularly against pathogenic Gram-negative bacteria in clinical and agricultural settings. Successful phage use has been demonstrated against Gram-negative rods and other bacterial pathogens in compassionate-use cases^6–8^, recent and ongoing clinical trials ^9,10^, and in agricultural pathogen control^11^.

Purifying phages from inflammatory bacterial contaminants generated during production, especially the endotoxin lipopolysaccharides (LPS), and antigenic or virulent host proteins, is a critical step before their therapeutic or agricultural use and remains a bottleneck in the broader application of phages. Cell-free production processes do exist for producing some phages, notably T7^12–15^, eliminating the need for extensive purification, but most phages are still produced in bacterial hosts^16^. Beyond ensuring safety when administered to humans and animals, phage purification is required for reproducible research in phage immunology and pharmacology, where presence of endotoxin can confound the interpretation of phage impacts on cells and animal models, as well as in studies of phage chemistry and biophysics^16–24^.

LPS is a potent pyrogen and trigger of innate immune activation. Concentrations as low as 0.025 EU/mL (∼0.0025 ng/mL) can induce detectable pro-inflammatory changes in monocytes in vitro^25,26^. Concentrations of 5–20 EU/kg can cause mild systemic symptoms in humans^27^, while higher concentrations can lead to septic shock and death^28,29^. The U.S. Food and Drug Administration (FDA) limits endotoxin levels in injectable drug products to < 5 EU kg^−1^ h^−130^. For agricultural applications, many products target ≤ 25,000 EU mL^−1^ to align with FDA “Generally Recognized as Safe” (GRAS) standards for food additives.

Host cell proteins (HCP) such as bacterial proteins are foreign molecule to the human body and might cause immune responses, cross-reaction with human proteins or autoimmunity effects^31,32^ . The interference with physiological processes creates a safety risk in phage therapy due to phage production within bacterial hosts^33^. Therefore, a close monitoring of low HCP levels is crucial to control for product quality in phage therapy^34^.

Monolithic chromatography is well suited for purifying phages and other viruses. Its large, continuous channels promote convective, low-shear flow, unlike porous beads or membranes, which generate turbulent, high-shear flow ^35,36^. Monoliths operate efficiently at high flow rates and across a range of particle sizes and viscosities, with reduced pressure drop and improved virion recovery^37^. Their surfaces can also be functionalized with diverse solid-phase chemistries, including preferential-exclusion, ion-exchange, and hydrogen-bonding ^38,39^. In preferential-exclusion chromatography, hydrophobic interactions between phages and the column wall are used to separate phages from LPS and other contaminants. Kosmotropic salts (e.g., potassium phosphate) are excluded from the hydration layer around biomolecules (phages) due to favorable bulk water–salt interactions. On hydroxyl-substituted (–OH) monolithic media, there is similarly an ion-depleted layer at high kosmotropic salt concentrations, which creates a hydrophobic attraction between phages and the column wall. As phages adsorb, the salt-free area and free energy decrease, further promoting adsorption. Reducing the salt concentration disrupts these interactions, weakening hydrophobic binding and eluting phages off the column^39–41^. In this study, we use preferential-exclusion chemistry in OH-monolithic column to purify phages

Prior work using monolithic columns shows capture and purification across formats but often requires multiple steps. Quaternary amine (QA) and diethyl aminoethyl (DEAE) anion-exchange monolithic columns can bind morphologically diverse phages (with recovery spanning 40-99%), though LPS removal has been limited, inconsistent, and/or not reported^42–44^. Purification of filamentous M13 using an OH-monolithic column and steric-exclusion chromatography of has been reported, with high recovery (90%) and host cell protein reduction (98%)^38^, but this has not been demonstrated for tailed phages. Two-step methods, such as OH-capture followed by polishing on a QA, H-bond, or PrimaS column, have achieved up to 7-log endotoxin reduction in LPS^39^.

Here, we report a single-step method utilizing monolithic OH-column chromatography and preferential-exclusion chemistry to purify multiple tailed phages with diverse physical characteristics. Using a 1 mL CIMmultus monolithic OH column and a potassium phosphate gradient, we purified five *Pseudomonas aeruginosa* phages (PAML-31-1, OMKO1, LPS5, TIVP-H6, DMS3vir), systemically comparing post-purification contaminant levels and recovery rates. Additionally, we linked recovery rates to size characteristics across both usual and a so-called “jumbo” phage^45,46^.

## Material and Methods

### Generation of clarified phage lysates

Bacteriophages were produced as described previously^20^. In brief, bacteriophages were propagated in a liquid culture in Erlenmeyer flasks. LB medium was inoculated with the *P. aeruginosa* host strain and incubated until an early to mid-log growth phase was reached (37 °C, 275 x *rpm*). Planktonic cultures were infected with phages at a multiplicity of infection (MOI) of 0.01 and incubated overnight. Gross bacterial debris was removed by two to three cycles of centrifugation (8000 x *g*, 20min 4°C) and the supernatant was filtered through a 0.22 µm polyethersulfone (PES) membrane (Corning, NY, Product No. 4311188). The lysate was treated with 0.05 U/mL Benzonase Nuclease (Sigma-Aldrich, St. Louis, MO, USA; Cat. No. E8263) overnight to digest free DNA.

### Monolithic chromatography phage purification

Benzonase-treated phage lysate was then mixed with 3 M KH_2_PO_4_ (potassium phosphate) buffer (pH 7.0) at a 1:1 ratio (to give a final concentration of 1.5 M KH_2_PO_4_, in a final volume of 10-60mL), in preparation for purification using a fast protein liquid chromatography (FPLC) instrument equipped with a P960 sample pump (ÄKTA Purifier, GE Healthcare Biosciences, Sweden. Purification was conducted with a CIMmultus OH 1 mL monolithic column with a 2 μm channel size (Sartorius BIA Separations, Ajdovščina, Slovenia). Column washing, regeneration, performance test, and storage were performed according to the manufacturer’s recommendations. Chromatography runs were analyzed with UNICORN 5.0 software (Cytiva Life Sciences, Marlborough, MA, USA). All purification protocols were run at room temperature, and all buffers and samples were filtered through a 0.22 µm PES membrane prior to use to remove particulates. For each run 10-60 mL of phage lysate in 1.5 M KH_2_PO_4_ was loaded onto a column pre-equilibrated with 1.5 M KH_2_PO_4_ loading buffer, pH 7.0. After sample application, the column was washed with 10 column volumes (CV) of loading buffer to remove unbound particles. Phages were eluted from the column using a linear gradient from 1.5M to 20 mM KH_2_PO_4_ over 20 column volumes. The fraction corresponding to the eluted phage was collected using a fraction collector based on inline measurement of UV absorbance at 280 nm by the FPLC instrument. Between purification of different phages, the column was cleaned by running a 1M NaOH solution through the column for 4 h at 0,2mL/min, followed by the manufacturers recommended clean-in-place procedure, which involves 1M NaOH cleaning, followed by a pH neutralization step with 200 mM Tris pH 7.5 and then a wash with ddH_2_O, before re-equilibration with 1.5 M KH_2_PO_4_ loading buffer, pH 7.0.^47^ See protocol.io: dx.doi.org/10.17504/protocols.io.5jyl8xn69v2w/v1 for details. Private link for reviewers: https://www.protocols.io/private/BDC73844E27F11F0BD000A58A9FEAC02 to be removed before publication.)

### Phage enumeration

Phages were quantified as described previously^20^. Plaque assays using the spot dilution double agar overlay method were employed. In brief, 100 μL of mid-log phase bacteria was added to molten 5 mL of top agar (5 g/L agar, 10 g/L tryptone, 10 g/L NaCl, 20 mM MgSO_4_, 20 mM CaCl_2_) held at 55 C, mixed by inversion, poured evenly onto the surface of a square 1.5% LB-agar plate, and allowed to solidify. 10-fold serial dilutions of phages were prepared in 96-well polystyrene U-bottom plates (ThermoFisher Scientific, Waltham, MA, USA, Cat. No. 168136) in SM+gel buffer (50 mM Tris-HCl, 100 mM NaCl, 8 mM MgSO_4_, 0.01% gelatin (Sigma-Aldrich, St. Louis, MO, USA, Cat. No. G7041), pH 7.50). 10 μL of each dilution was spotted onto the top agar. Spots were allowed to fully dry with lids open in a biosafety cabinet, prior to plates being inverted and incubated at 37 °C overnight. Visible plaques were counted and used to calculate the phage titer in PFU/mL.

### Endotoxin, protein and DNA Quantification

The endotoxin level of samples was measured using a recombinant Factor C Endotoxin Detection assay (bioMérieux, France, Endonext EndoZyme II assay Cat. No. 423129) with an assay range from 0.005 to 50 endotoxin units (EU)/mL. Samples were diluted with endotoxin-free water to attain the assay range according to the manufacturer’s instructions. The total protein content of samples was measured using a bicinchoninic acid (BCA) assay (Thermo Scientific, USA, Pierce BCA Protein Assay kit Cat. No. 23225) according to the manufacturer’s instructions. The total DNA was determined by a Quant-iT PicoGreen assay (Invitrogen, USA Cat. No. P11496) Endotoxin, protein and DNA readings were performed in 96-well plates using an automated, multifunctional microplate reader (Tecan SPARK multimode reader, Switzerland).

### Transmission electron microscopy (TEM)

TEM imaging was done as previously reported^19^. In brief, the size and morphology of phages were examined using a JEOL JEM1400 transmission electron microscopy (TEM) (JEOL USA Inc., Peabody, MA) at 80 kV. 5 µL of diluted phage solution was dropped onto a carbon-coated copper grids (FCF-200-Cu, Electron Microscopy Sciences, Hatfield, PA). After 3 minutes, the grid was dipped into a ddH_2_O droplet and laid sample side up. Then 1% uranyl acetate for staining was dropped onto the sample and was allowed to dry for 15 minutes before performing microscopy.

### Dynamic light scattering (DLS)

DLS measurement was done as previously reported^19^. All DLS size distributions were conducted with a Nano-Series Zeta Sizer (Nano-NS ZEN3600; Malvern Instruments, Worcestershire, UK) equipped with a 633 nm laser. Measurements were obtained at 25 °C at a backscattering angle of 173°. Each DLS measurement reported in this study is the average of three replicates. Each replicate is itself an average of eleven 10 s measurements obtained after a 1 min vortexing period on half-speed, 1 min equilibration period in the instrument, and with variable attenuation to generate count rates >100 kcps. Phage diffusion coefficients were calculated from auto-correlated light intensity data, and hydrodynamic diameters (*D*_H_) were computed using ZetaSizer version 7 software with the Stokes-Einstein equation.

## Results

### Single-step OH-monolithic chromatography can purify phages with diverse physical characteristics, maintaining structural activity and bioactivity

We purified five different *Pseudomonas aeruginosa* phages on a FPLC instrument equipped with a CIMmultus 1 mL OH-monolithic column (**Fig. 2**). All were tailed *Caudoviricetes* phages: four myoviruses (PAML-31-1, LPS-5, OMKO1) and two siphoviruses (TIVP-H6 and DMS3vir). TEM showed PAML-31-1 (head length 75 +/- 2 nm; tail length 147 +/- 5 nm) and LPS-5 (head length 76 +/- 2 nm; tail length 143 +/- 4 nm) were the smallest and of similar dimensions, while OMKO1 was substantially larger (head length 129 +/- 9 nm; tail length 200 +/- 19nm; “jumbo phage” (**Fig. 2A**). TIVP-H6 (head length 68 +/- 4 nm; tail length 178 +/- 12 nm) displayed the expected long, flexible siphovirus tail; DMS3vir had a smaller head and longer tail (head 44 +/- 2 nm; tail 225 +/- 1 nm).

Post-purification DLS **(Fig. 2B)** showed a single peak in the size-intensity spectrum for each phage with low polydispersity (PDI) (PDI = 0.093-0.188) and Z-averages consistent with the TEM-inferred sizes (e.g. PAML-31-1 94 nm; LPS-5 147.6 nm; OMKO1 218.1 nm; TIVP-H6 113 nm; DMS3vir 122.4 nm). Together with the infectivity measurements by plaque assays **(Table 1)**, these data indicate that OH-monolith chromatography yields active, uniform, monodisperse phage preparations. Additionally, the data show that the elution buffer conditions effectively eluted phages without compromising the structural integrity of the phages, and that the clean-in-place protocol eliminated cross-contamination of phages between runs.

**Table 1:**
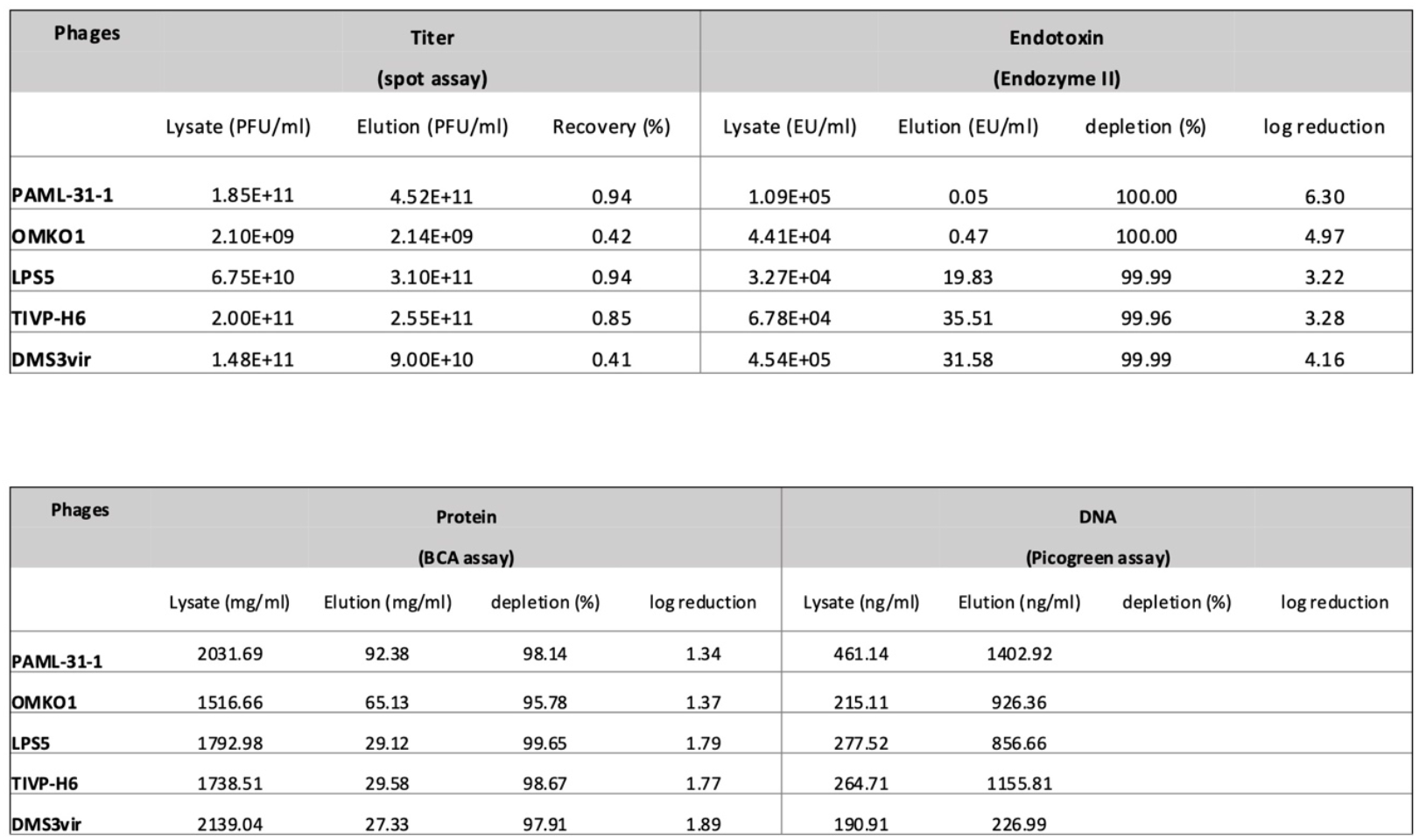
Summary of purification results. Titer (plaque assay), endotoxin (Endozyme II assay), protein (BCA assay), and DNA (Picogreen assay) concentrations before and after chromatography using the CIMmultus OH-column of five lytic Pseudomonas aeruginosa phages. Calculated values are phage recovery (%), impurity depletion (%), and impurity log reduction.

### OH-monolithic chromatography demonstrates high recovery for myoviruses and siphoviruses

Clarified lysates of the five *P. aeruginosa* phages (PAML-31-1, LPS-5, TIVP-H6, OMKO1, DMS3vir) were adjusted with concentrated 3 M KH_2_PO_4_ (pH 7.0) to 1.5 M final concentration (to achieve same concentration of KH_2_PO_4_ as the binding/equilibration buffer), loaded on an equilibrate CIMmultus 1 mL OH column at 1 mL min^-1^, and eluted by a linear decrease from 1.5 M to 20 mM KH_2_PO_4_ over 20 CV **(Fig. 1)**. A representative chromatogram **(Fig. 1F)** shows high UV absorbance at 280 nm during the loading step, reflecting proteins and other factors absorbing at 280 nm in the lysate that did not bind the column and a single sharp peak during the elution phase, where proteins that bound to the column were eluted **(Fig. 1, chromatogram)**. The sharp peak indicates a uniform population. Plaque assays of pooled fractions corresponding to the peak demonstrated high recoveries for PAML-31-1 and LPS-5 (both 94%; elution titers 4.52 × 10^11^ and 3.10 × 10^11^ PFU mL^-1^ respectively), good recovery for TIVP-H6 (85%; 2.55 × 10^11^ PFU mL^-1^), and lower recoveries for DMS3vir (41%; 9.00 × 10^10^ PFU mL^-1^) and the jumbo phage OMKO1 (42%; 2.14 × 10^9^ PFU mL^-1^) **(Table 1)**. Interestingly, we observed a column pressure increase during sample application, for the jumbo phage OMKO1 relative to the smaller LPS-5, loading an identical volume of 40 mL of each phage lysate. (**Fig. 1 supplementary data**)

**Figure 1:**
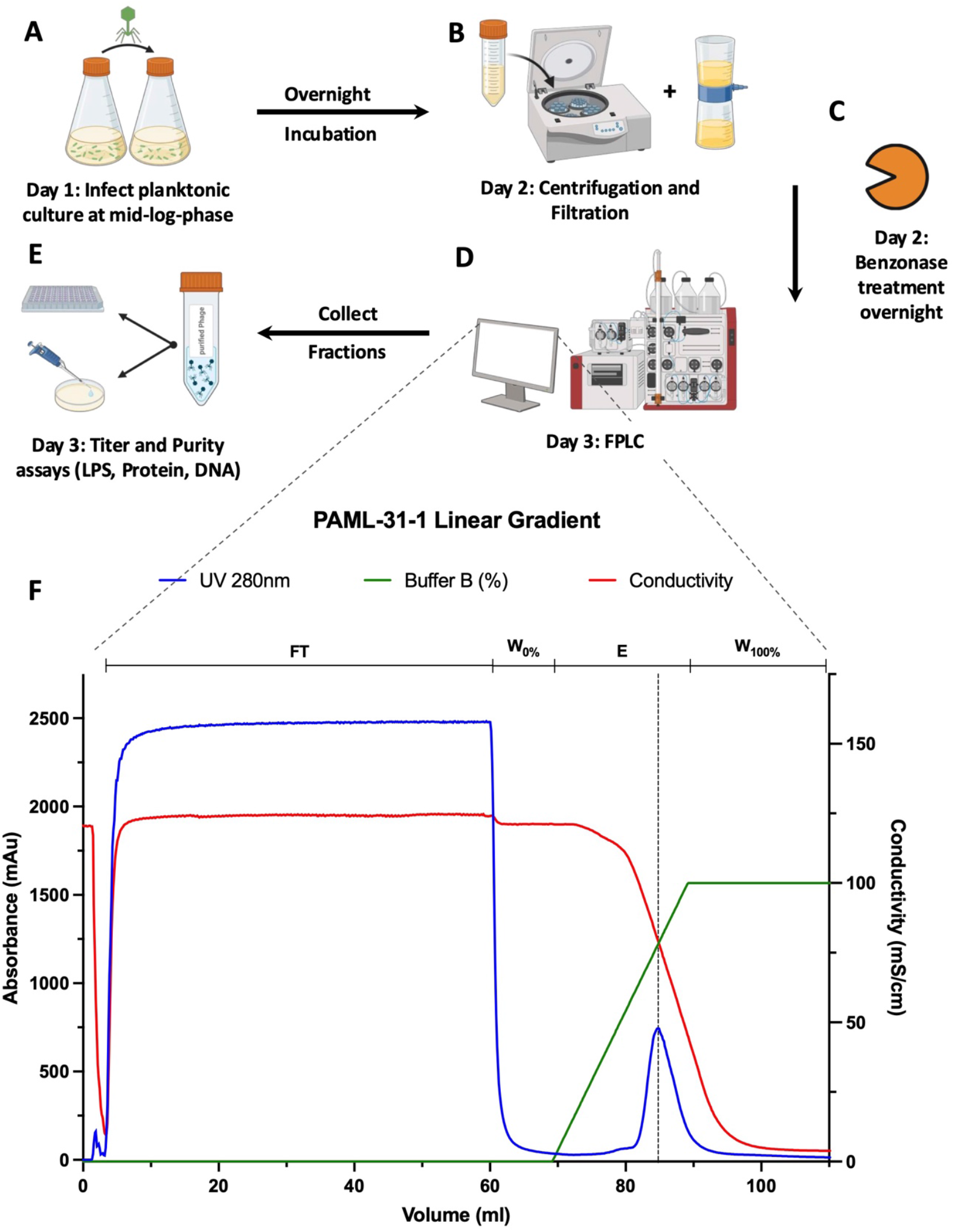
Monolithic Chromatography preparation of PAML-31-1 phage clarified lysate on CIMmultus OH 1ml column on AKTA purifier FPLC system. 60 ml of PAML-31-1 phage lysate diluted to 1.5 M KH_2_PO_4_, pH 7.0 is loaded. Loading Buffer A: 1.5 M KH_2_PO_4_, pH 7.0, Elution Buffer B: 20mM KH_2_PO_4_, pH 7.0. Upper lines indicate different phases of the method: Flowthrough phase (**FT**T), 10 CV washing the column after sample application with 10 CV (**W**_**0%**_), Elution with a linear Gradient over 20 CV to 100% Elution Buffer (**E**), final wash with 100% Buffer B to regenerate the column (**W**_**100%**_). Detection: blue: UV-absorbance (280 nm), red: Conductivity (mS/cm), green: Buffer B (%). Created in BioRender.com

**Figure 2:**
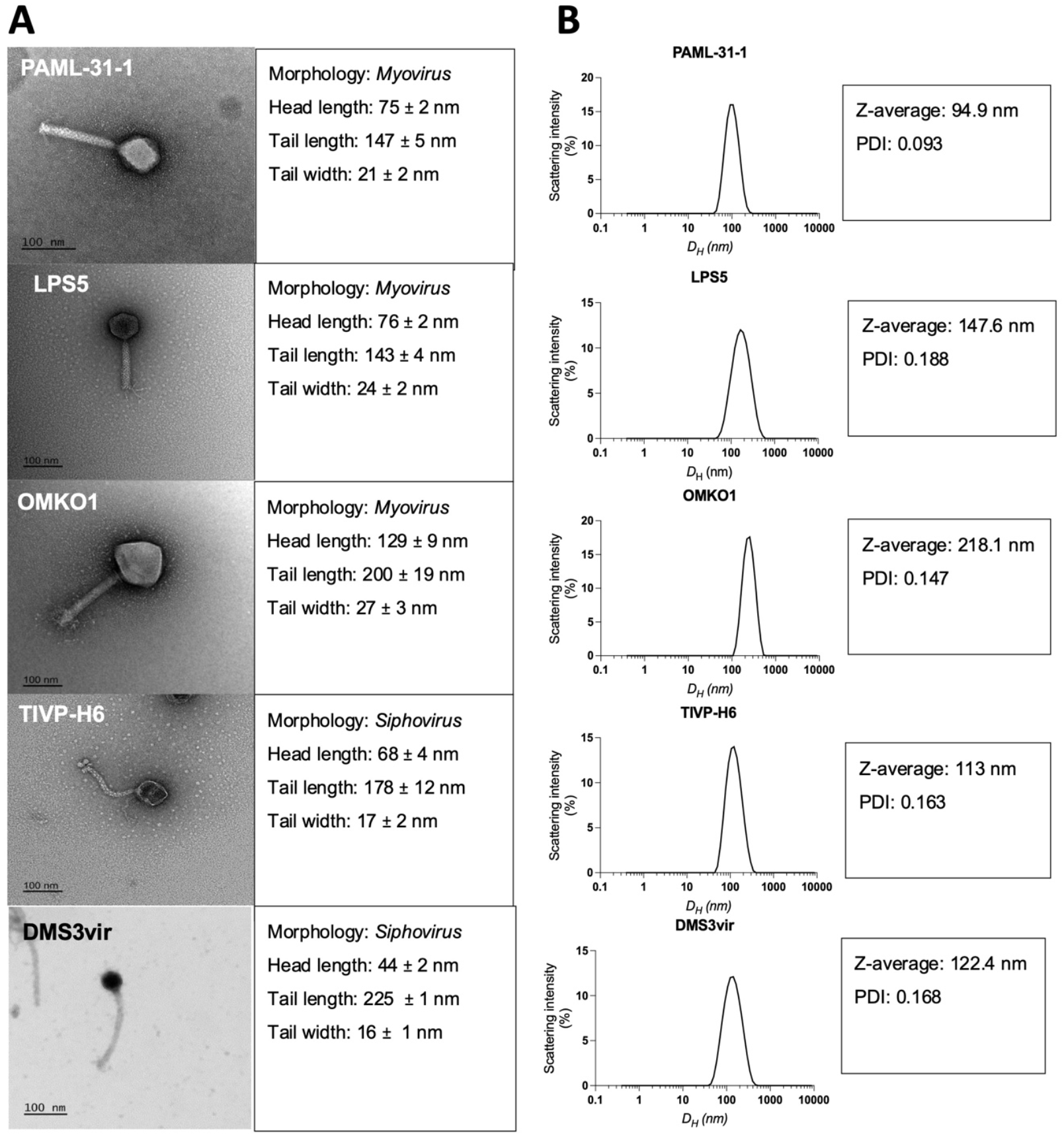
Size characteristics of 5 lytic *P. aeruginosa* phages used in this study. **A**. A representative transmission electron microscopy image of every is shown at 80,000x magnification. Head length, tail length, and tail width are measured on at least 5 different transmission electron microscopy examples for each phage. **B**. Dynamic light scattering (DLS) spectra, Z-average and PDI values of purified phages used in this study.

### OH-monolithic chromatography substantially reduces endotoxin and host protein in clarified P. aeruginosa phage lysates

After debris removal by centrifugation and filtration, lysates still contained high impurity loads (lysate Endozyme II endotoxin measurement ∼1.1-4.5 × 10^5^ EU mL^-1^; lysate BCA protein measurement ∼1.5-2.1 g mL^-1^). Following a 10-CV wash and a 20-CV linear gradient elution, endotoxin in the pooled infectious fractions was depleted by 99.96-100.00% across the five phages log_10_ reduction 3.22-6.30) e.g., PAML-31-1 to 0.05 EU mL^-1^ (6.30-log, 100%), OMKO1 to 0.47 EU mL^-1^ (4.97-log, 100%), LPS-5 to 19.83 EU mL^-1^ (3.22-log, 99.99%), TIVP-H6 to 35.51 EU mL^-1^ (3.28-log 99.96), and DMS3vir to 31.58 EU mL^-1^ (4.16 log 99.99%). Protein was considerably reduced in the same fractions, with 95.8-99.7% depletion (log_10_ reduction 1.34-1.89): e.g., PAML-31-1 to 92.38 mg mL^-1^ (1.34-log, 98.14%), OMKO1 to 65.13 mg mL^-1^ (1.37-log, 95.78%) LPS-5 to 29.12 mg mL^-1^ (1.79-log, 99.65%), TIVP-H6 to 29.58 mg mL^-1^ (1.77-log, 98.67%), and DMS3vir to 27.33 mg mL^-1^ (1.89-log, 97,91%) **(Table 1)**. By contrast, we observed DNA levels, as measured by PicoGreen, to increase rather than decrease in the eluted phage fractions (from a mean of 282 ng mL^-1^ in lysates to 914 ng mL^-1^ post -purification across the five phages).

### Recovery is partly explained by transport constraints in convective monoliths

To test whether particle size explains the observed difference in recovery, we analyzed mass transfer in convective monoliths, where pore diffusion is negligible and film resistance at the column wall dominates. The convective mass-transfer coefficient k_f_ (m s^-1^) where Sherwood number is represented as (Sh), free-solution mass diffusivity D_0_ (m^2^ s^-1^), and characteristic length L (m).

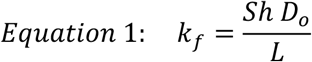

Diffusivity was estimated from DLS-derived hydrodynamic diameters using the Stokes-Einstein equation, where Boltzmann’s constant is represented as k_B_, temperature T, dynamic viscosity η, and hydrodynamic diameter D_H_.^48^

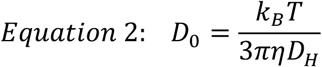

with Boltzmann constant k_B_, temperature T, dynamic viscosity η, and hydrodynamic diameter D_H_.

At k_B_ = 1.38 × 10^−23^ J K^−1^, T = 298.15 K, η = 8.9 × 10^−4^ Pa s, and using the hydrodynamic diameters from the DLS measurements, we obtained mass diffusivity (D_0_) for PAML-31-1 (D_0_ = 5.17 × 10^-21^ m^2^ s^-1^) and OMKO1 (D_0_ = 2.25 × 10^-21^ m^2^ s^-1^) The diffusivity ratio of 0.435 closely corresponded to the observed relative recovery fraction of 0.435, consistent with film-transfer contributions to observed yield differences. Across five phages, recovery decreased with hydrodynamic diameter **(Fig. 3B)** and observed recovery correlated with expected recovery (Pearson r = 0.552, r^2^ = 0.305; **Fig. 3C**), indicating that diffusivity accounts for ∼30% of the observed variance in recovery.

**Figure 3.**
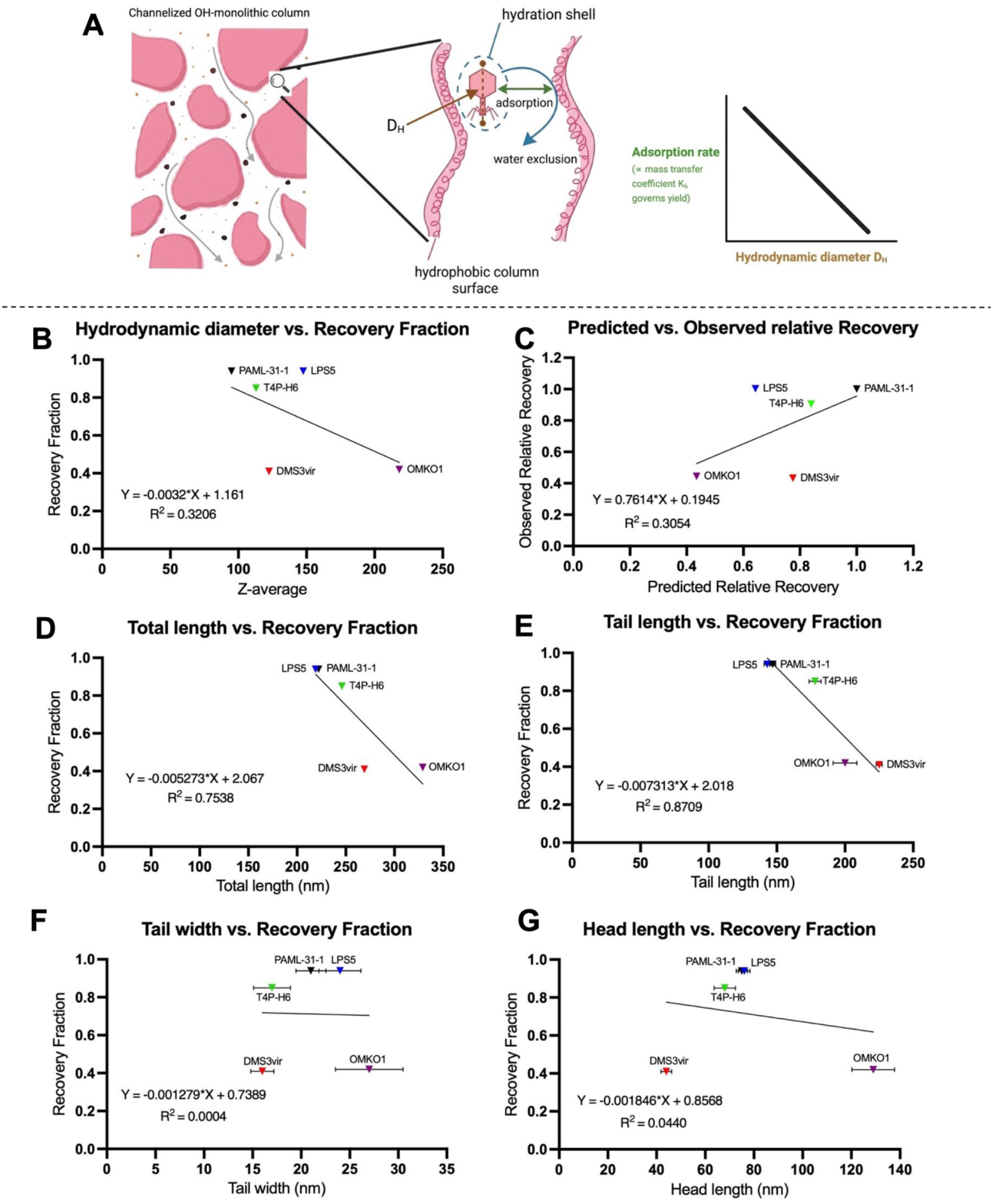
Size–transport relationships governing phage recovery on OH monoliths. **(A)** Schematic of convective mass transfer in a monolithic channel: bulk flow bypasses pores (no intraparticle diffusion). Phages cross a thin interfacial film at the wall bearing OH ligands; the mass-transfer coefficient *k*_*f*_ scales with free-solution diffusivity ***D***_**0**_ per Eq. 1, and *D*_**0**_ is inversely proportional to hydrodynamic diameter ***D***_***H***_ per Eq. 2, Stokes–Einstein. **(B)** Hydrodynamic diameter **(*D***_***H***_, DLS Z-average) vs. Recovery fraction. **(C)** Observed recovery vs predicted recovery proportional to *D*_**0**_ computed from Eq. 2. **D)** Total virion length (head + tail) vs. Recovery; negative trend partially collinear with ***D***_***H***_. **(E)** Tail length vs. Recovery; strong negative associatio**n. (F)** Tail width vs. Recovery; no association. **(G)** Head length vs. Recovery; no association. Points denote individual phages (PAML-31-1, LPS-5, TIVP-H6, DMS3vir, OMKO1). Error bars indicate SD. Created in BioRender.com

We also noted that recovery fraction fell sharply with increasing tail length and total virion length (both dominated by tail length), with long-tailed phages such as DMS3vir showing low recovery despite only moderate hydrodynamic diameter **(Fig. 3D)**. Head length and width showed little association. During sample application, column pressure increased for OMKO1 relative to LPS-5 (**Fig. 1 supplementary data**. Together, these observations suggest that phage recovery from the column is primarily limited by size-dependent mass transfer but can also be impacted by extreme tail length and morphology.

## Discussion

We show that OH-monolithic chromatography on CIMmultus columns provides a rapid, reliable, single-step capture strategy for tailed *P. aeruginosa* phages (Myoviridae: PAML-31-1, LPS-5, OMKO1, DMS3vir; Siphoviridae: TIVP-H6). Recoveries were high for PAML-31-1, LPS-5, and TIVP-H6, but lower for OMKO1 and DMS3vir. Infectious eluates showed substantial reductions in host protein and endotoxin, reaching levels compatible with commonly cited limits and regulatory thresholds for intravenous delivery or ingestion at typical dosing, underscoring translational potential. For a typical intravenous phage dose of a typical intravenous phage dose of 10^9^ PFU delivered in 1 mL over 1 hour to a 70-kg patient, the endotoxin levels achieved here (0.05-35.51 EU/mL) translate to 0.0007-0.51 EU/kg, well below the FDA limit of 5 EU/kg/h.

Mechanistically, our data support a transport constraint arising from film-limited mass transfer in convective monoliths. Because pore diffusion is negligible, transfer to the stationary phase scales with the free-solution diffusivity D_0_, which is inversely related to the hydrodynamic diameter by Stokes-Einstein. This framework explains a substantial portion of the observed yield differences, shown earlier in the particular case of OMKO1 where differences in hydrodynamic diameter were large, but consistently true across phages with moderate positive correlation between observed and expected recovery. These data support routine DLS measurement of phage hydrodynamic diameters to inform purification strategies.

We also found a strong negative association between tail length and recovery and a similar but weaker trend with total length, whereas head length/width showed little association. Notably, DMS3vir showed low recovery despite small head size, indicating contributors beyond small head size alone, such as shear sensitivity, tail architecture, and virion surface chemistry may play an important role.

Operationally, this workflow is straightforward (∼3 days) and avoids potentially hazardous reagents (no CsCl, chloroform, or Triton X-100). The OH monolith tolerates 1 M NaOH clean-in-place between runs without detectable cross-contamination, supporting reuse and economical scale-up to larger monoliths. While we achieve therapeutic-grade purity with a single step, the process is modular and can be paired with alternative CIM chemistries when phage surface properties warrant polishing or when other specific impurities must be minimized.

An important analytical point is that increases in PicoGreen-measured DNA in infectious eluates should be interpreted as co-elution of encapsulated phage genomes with active particles rather than carry-through of free DNA. One limitation of this study is the lack of nuclease-resistant controls or capsid-permeabilization assays to prove the free DNA has been reduced.

Our findings extend prior single-phage demonstrations of OH-monolith capture (e.g., by Rebula *et al*. for PP-01) by showing that a single-column OH process can generalize across multiple *P. aeruginosa* phages while maintaining high recovery for most and delivering clinically relevant purity for all. Compared with workflows that rely on sequential column chemistries, this single-step approach reduces processing time, limits inter-step losses, and simplifies technology adoption.

Additional limitations include low recovery for OMKO1 and DMS3vir under the present conditions, small sample size (five phages), and reliance of DLS Z-averages for hydrodynamic diameters. Future work should test parameter levers predicted by the transport model such as load rate, gradient length/shape, temperature/viscosity, and column volume, and evaluate gentler binding/elution conditions to further mitigate shear. Phage-specific interactions could be addressed by tuning buffer composition and strength.

A single-step chromatographic purification on CIMmultus OH-monolithic columns efficiently generates therapeutic-grade phage preparations across multiple morphologies, delivering high recoveries for most phages and therapeutic-grade purity in just one run. This method is simple, scalable, and effective at endotoxin clearance and host protein removal, and the column reusability (NaOH clean-in-place) offers practical advantages for both research and production. Importantly, the size-transport analysis explains a significant proportion of variation in yield, and provides clear levers (flow, gradient shape/length, temperature/viscosity, and column scale) to optimize the purification of challenging virions (e.g., OMKO1, DMS3vir). Together, these features position OH-monolith capture as a robust platform for advancing phage purification for clinical and agricultural applications.

## Data availability

Data are available upon reasonable request from the corresponding authors subject to institutional review and approval. Supporting data values associated with the main manuscript is available in supplementary materials.

## Author Contributions

Conceptualization: PLB, FB, AE, TD. Methodology: AE, TD, MH, PB. Investigation and Data Processing: AE, TD, MH, PB, FB. Data Analysis: AE, TD, JCS, PLB. Writing: PLB, JCS, TD, AE.

## Acknowledgments

We gratefully acknowledge the following funding: National Institutes of Health grant R01 HL148184-01, R01 AI12492093, R01 DC019965, and grants from the Cystic Fibrosis Foundation (CFF), the Cystic Fibrosis Research Institute (CFRI) and the Emerson Collective grant (to PLB) and grants from the Stanford University Medical Scientist Training Program grant T32-GM007365 and the Stanford Interdisciplinary Graduate Fellowship, Gold Family Graduate Fellow (to TD) and the Merrigan Fellowship (to AE). This project was funded in part with Federal funds from the National Institute of Allergy and Infectious Diseases, National Institutes of Health, Department of Health and Human Services, under Contract No. HHSN272201700020I, Task Order 75N93021F00003 (A-57).

