## Supplemental data for "Single-step monolithic chromatography efficiently purifies diverse Pseudomonas aeruginosa phages with therapeutic-grade endotoxin reduction"

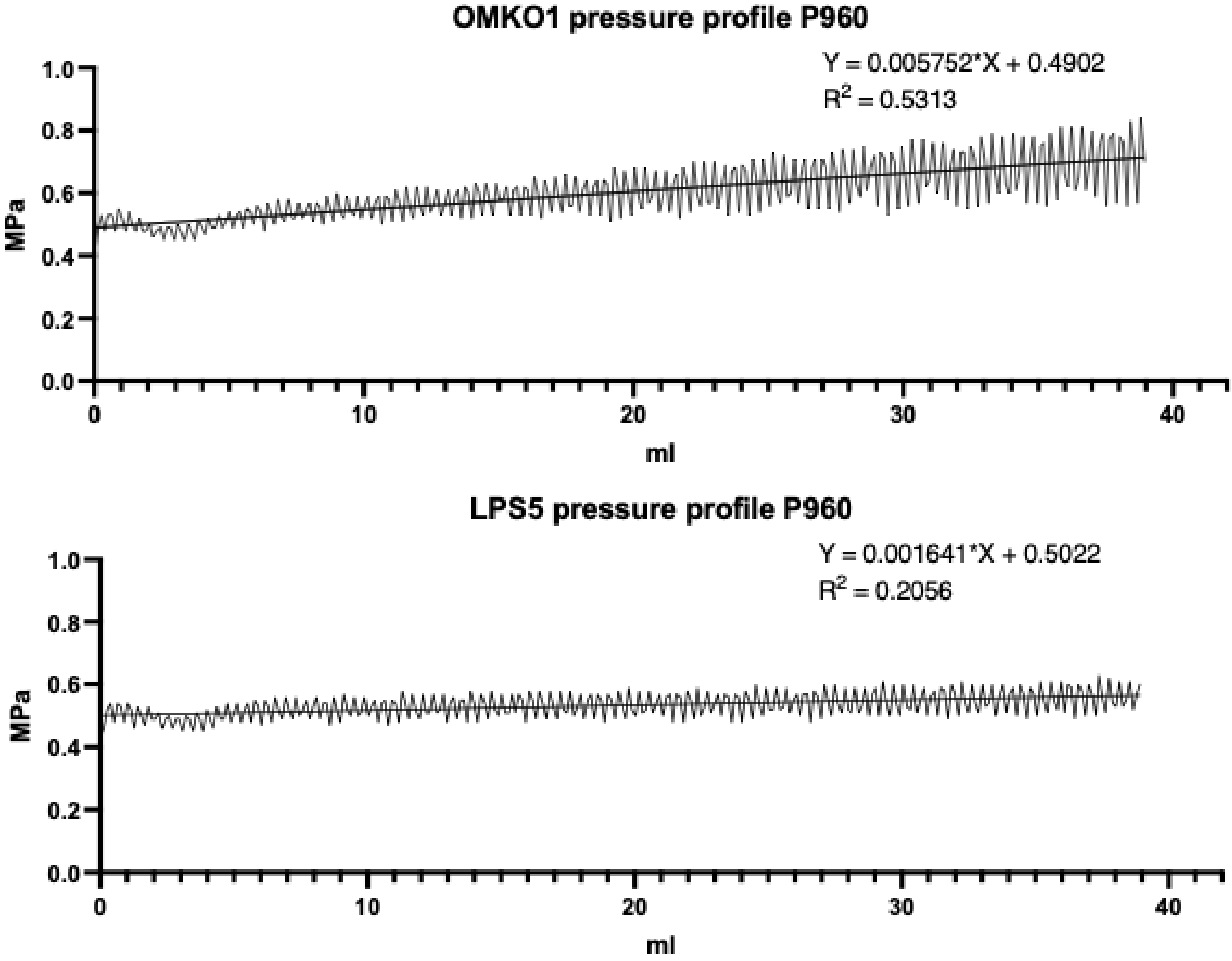


**Figure 1: Comparison of pressure increase during the sample loading phase between OMKO1 and LPS5.** 40 ml of OMKO1 and LPS5 phage lysates diluted to 1.5 M $KH_2PO_4$, pH 7.0 loading buffer conditions were loaded into CIMmultus OH 1ml column connected to an FPLC system with a sample pump P960. The pressure increase (MPa) within loading phase was measured by the P960 sample pump and plotted over volume (ml). A simple linear regression was performed on given data points. Equations and R2 are displayed.

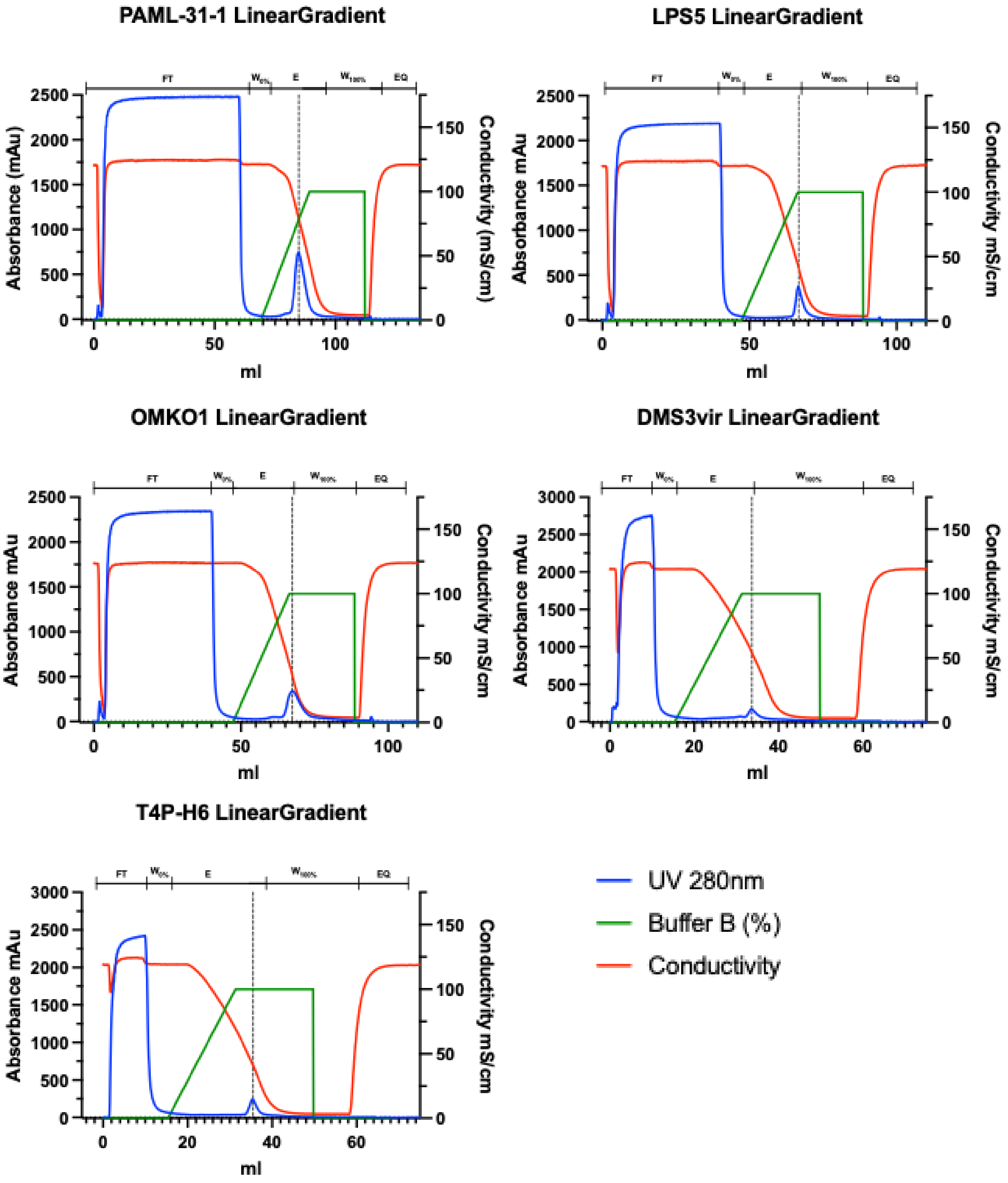


**Figure 2: Monolithic Chromatogram of five phage clarified lysates on CIMmultus OH 1ml column on Akta purifier FPLC system.** Distinct volumes of of PAML-31-1, LPS5, OMKO1, DMS3vir, T4P-H6 phage lysates diluted to 1.5 M $KH_2PO_4$, pH 7.0 is loaded. Loading Buffer A: 1.5 M $KH_2PO_4$, pH 7.0, Elution Buffer B: 20mM $KH_2PO_4$, pH 7.0. Upper lines indicate different phases of the method: Flowthrough phase (**FT**), 10 CV washing the column after sample application with 10 CV (**$W_{0\%}$**), Elution with a linear Gradient over 20 CV to 100% Elution Buffer (**E**), final wash with 100% Buffer B to regenerate the column (**$W_{100\%}$**). Detection: blue: UV-absorbance (280 nm), red: Conductivity (mS/cm), green: Buffer B (%).

**PAML-31-1**

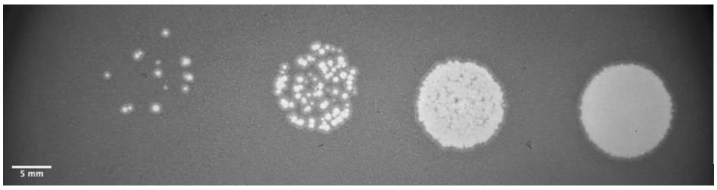


Morphology: Clear, well circumsribed

Plaque diameter: 0.915 ± 0.379 mm

**LPS5**

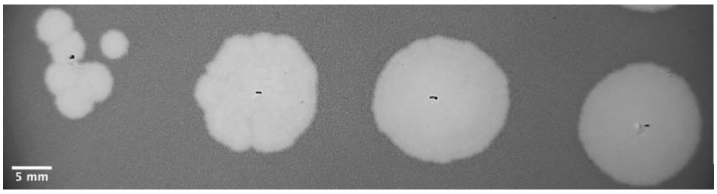


Morphology: Clear, well circumsribed

Plaque diameter: 4.31 ± 0.304 mm

**OMKO1**

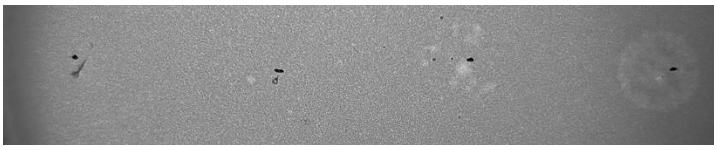

Morphology: faint

Plaque diameter: 1.12 ± 0.236 mm

**DMS3vir**

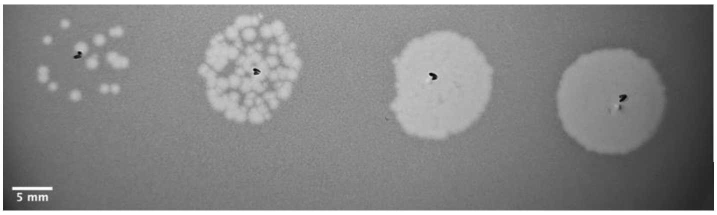


Morphology: Clear, well circumsribed

Plaque diameter: 1.48 ± 0.276 mm

**T4P-H6**

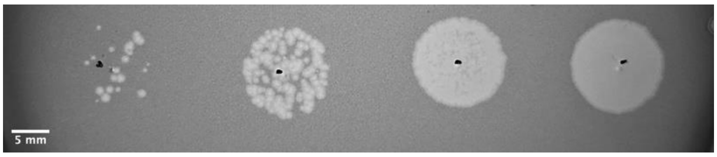


Morphology: Clear, well circumsribed

Plaque diameter: 1.09 ± 0.468 mm

**Figure 3: Plaque morphology and plaque diameter of five phages used in this study.** Demonstration of different plaque morphologies across the five used phages (PAML-31-1, LPS5, OMKO1, DMS3vir, T4P-H6). Plaque assays were conducted using the spot dilution double agar overlay method. 10 µL of 10-fold serial dilutions was spotted onto the top agar. After incubation, Visible plaques were counted and used to calculate the phage titer in PFU/mL. Different dilutions were imaged depending where plaques were most distinguishable.
